# MITOMIX, an Algorithm to Reconstruct Population Admixture Histories Indicates Ancient European Ancestry of Modern Hungarians

**DOI:** 10.1101/247395

**Authors:** Zoltán Maróti, Tibor Török, Endre Neparáczki, István Raskó, István Nagy, Miklós Maróti, Tamás Varga, Péter Bihari, Zsolt Boldogkői, Dóra Tombácz, Tibor Kalmár

## Abstract

By making use of the increasing number of available mitogenomes we propose a novel population genetic distance metric, named Shared Haplogroup Distance (SHD). Unlike F_ST_, SHD is a true mathematical distance that complies with all metric axioms, which enables our new algorithm (MITOMIX) to detect population-level admixture based on SHD minimum optimization. In order to demonstrate the effectiveness of our methodology we analyzed the relation of 62 modern and 25 ancient Eurasian human populations, and compared our results with the most widely used F_ST_ calculation. We also sequenced and performed an in-depth analysis of 272 modern Hungarian mtDNA genomes to shed light on the genetic composition of modern Hungarians. MITOMIX analysis showed that in general admixture occurred between neighboring populations, but in some cases it also indicated admixture with migrating populations. SHD and MITOMIX analysis comply with known genetic data and shows that in case of closely related and/or admixing populations, SHD gives more realistic results and provides better resolution than F_ST_. Our results suggest that the majority of modern Hungarian maternal lineages have Late Neolith/Bronze Age European origins (partially shared also with modern Danish, Belgian/Dutch and Basque populations), and a smaller fraction originates from surrounding (Serbian, Croatian, Slovakian, Romanian) populations. However only a minor genetic contribution (<3%) was identified from the IX^th^ Hungarian Conquerors whom are deemed to have brought Hungarians to the Carpathian Basin. Our analysis shows that SHD and MITOMIX can augment previous methods by providing novel insights into past population processes.

## Background

In the era of Next Generation sequencing (NGS) an increasing amount of complete human mitochondrial genomes are available in public databases from both recent and archaic samples. Analysis of mtDNA genomes allows tracking of human evolution and migration as they refer to direct descent of the maternal lineages [1]. Based on mutational distances between individual mtDNA sequences their evolutionary genealogy have been reconstructed resulting in a phylogenetic tree [2]. The major branch points of the tree define mtDNA haplogroups (mt Hgs) while unique nucleotide combinations of individual mtDNA sequences are referred as haplotypes. The spatial distribution of mitochondrial haplogroups and haplotypes via human migrations shows distinct patterns, leading to the foundation of phylogeography [3].

The evolution of new mutations on the mitochondrial phylogenetic tree and their subsequent spreading takes many generations. Each new mutation in the mtDNA genome is prone to fixation, thereby giving rise to a new sub-branch on the phylotree, which can only happen at the geographical region where its parent mtDNA haplotype had already been prevalent. Because of geographical and cultural barriers the intra-population genetic mixture is obviously greater than inter-population genetic mixture, thus the emergence of new mtDNA haplogroups are initially always population specific, and unless migration occurs, they are confined to distinct geo-locations. The most recent mtDNA sub-haplogroups, appearing as “leaves” on the mitochondrial tree are necessarily less frequent then their progenitor lineages, and thus they are more population specific and provide even better geo-location due to the shorter duration of time available for their migration and admixture. Nevertheless population genetic processes like selection and genetic drift may eventually spread younger lineages at the expense of their progenitors. As older mt Hgs arose thousands of years earlier, they can be present on a more extensive geographical range due to migration and admixture; consequently, they are usually less localization- and population- specific in comparison to their descendant lineages. Despite their more extensive spreading, even older mt Hgs show a good correlation with larger geographical areas (continents), which is well-illustrated by the phylogenetic confirmation of the “Out of Africa” model [4, 5]. Experimental data indicate that fixation of new mutations is a very slow process, often requiring thousands of years [6] whereas on this timescale human migration and admixture can be considerable. For example the Migration Period of Europe in the middle of the first millennium AD or the colonization of America by White and Afro-American people happened in such a short time span, during which mtDNA sequence evolution is negligible. Thus considering the mitochondrial clock for such short periods of time, the mt Hg distribution of populations primarily reflects their admixture history and co-ancestry.

As human populations are seldom isolated, cultural exchanges regularly bring about admixture of diverse lineages, which is predominant between neighboring groups. As local admixture is a persistent process, on a larger time scale it partially conflates neighboring groups genetically, and could have a major effect on population structure. As a result, migrations deliver a set of locally admixed mt Hgs, characteristic for the donor population, to new locations. Thus most modern and archaic populations consist of individuals belonging to mt Hgs of various origins as also illustrated by our mitogenome database (Supplementary Table 1). According to available data approximately 5400 mt Hgs exist in archaic/recent populations [2], and each of these have a unique history concerning their geographical origin and spread in time and space [6]. Considering the emergence of thousands of mt Hgs at different times and locations on the two dimensional surface of the Earth and their independent dynamics, each geo-location is expected to have a unique mt Hg frequency distribution. For the same reason each population can be characterized by a unique subset of the currently known few thousand mt Hgs carried by their individuals in a population specific distribution. It can be postulated that this unique distribution could be used to track population histories in space and time, if sufficient data are available from different periods and locations. As all mt Hgs originated from a single progenitor, shared mt Hgs between two populations can only stem from common ancestors or population admixture, and the ratio of shared mt Hg lineages in admixing populations can be indicative of admixture ratios. Based on these considerations we introduce a new method for calculating population genetic distance called Shared Haplogroup Distance (SHD). From the mt Hg distribution of admixed populations their admixture history can be theoretically reconstructed with our new algorithm called MITOMIX, which estimates the best hypothetical admixture ratios based on SHD global minimum optimization in order to explain the observed mt Hg distributions of any studied population.

In order to highlight the usefulness of our newly proposed method we compared SHD and F_ST_ using our compiled database consisting of 62 modern and 25 archaic Eurasian human populations from which full length mt genome is currently available. Furthermore, we provide a detailed analysis of the genetic makeup of the modern Hungarians using 272 modern Hungarian mtDNA genomes.

## Results

### SHD calculation from mitochondrial haplogroup frequencies

The mitochondria can be viewed as a single polymorphic haploid locus which is inherited without recombination from mother to offspring. Due to the genesis and population dynamics of mt Hgs in space and time each of the geo-locations and populations can be characterized by a unique subset and distribution of mt Hgs and that shared mt Hgs indicate shared population history.

Let us assume that there are *n* mt Hgs. Let *x_i_*, and *y_i_* denote the frequency of the *i-th* haplogroup in populations *x* and *y*, respectively. We will identify the populations with their mt Hg distributions, thus the two populations will be denoted by *x*: = (*x*_1_, *x*_2_, …, *x_n_*) and *y*: = (*y*_1_, *y*_2_, …, *y_n_*), respectively. We can regard the two populations as random variables, in which case the chance that a random individual selected from population belongs *x* to haplogroup *i* will be *x_i_* = *P*(*x* = *i*); the probability of the random variable *x* taking the value *i*.

To calculate the mitochondrial Shared Haplogroup Distance of two populations from the differences between their mt Hg frequency distributions we introduce the following formula:

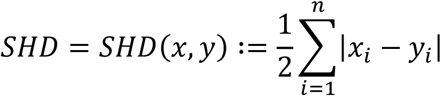

Note, that this is exactly the total variance distance of the two random variables (the two populations). Notice also that 0 ≤ *SHD* (*x*, *y*) ≤ 1 and *SHD* (*x*, *y*) = 0 if and only if *x*_*i*_ = *y*_*i*_ for all 0 ≤ *i* ≤ *n*. Moreover *SHD* (*x*, *y*) = 1 if and only if *x_i_* and *y_i_* are never simultaneously non-zero, that is *x_i_y_i_*= 0 holds for all 1 ≤ *i* ≤ *n*. In other words SHD is zero between two populations containing the same mt Hgs with identical frequencies, whereas it is 1 between two populations with no mt Hg overlap. Notice that the SHD formula introduced above satisfies the mathematical definition of a metric, as for any three (*x*, *y* and *z*) populations the following holds true:

1. *SHD* (*x*, *y*) = 0 if and only if *x* = *y*
2. *SHD* (*x*, *y*) = *SHD*(*y*, *x*)
3. *SHD* (*x*, *z*) ≤ *SHD* (*x*, *y*) + *SHD*(*y*, *z*)

The first and second are the trivial consequence of the formula 
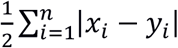
. Proving the third we need to show that for any three *x*, *y* and *z* probability distributions the triangle inequality holds true. Since taking the absolute value of the difference is the distance metric in one dimension, we have
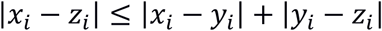
 for all *i*, and equality holds only in the case when *y_i_* lies in between *x_i_* and *z_i_*. From this we get

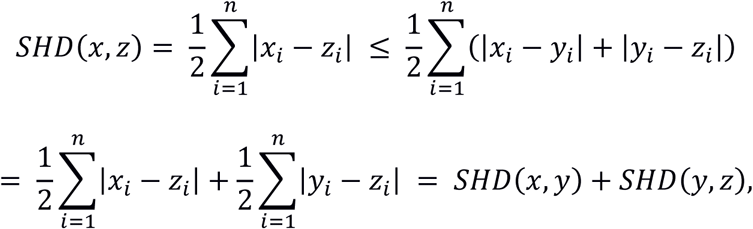

which concludes the proof. Thus the SHD value is the shortest path between two probability distributions in this metric space.

It can also be proven that for an arbitrary mixture between any two populations resulting in an admixed population, the SHD values between the parents and the resulting admixed population are proportional to the ratio of parents in the mixture.

#### Proposition

let *x* and *y* be the probability distributions of parent populations, *z* the distribution of the admixed population and *ε* the mixing ratio, thus *z* = *εx* + (1 − *ε*)*y*. Then

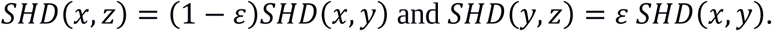

In particular *SHD* (*x*, *y*) = *SHD* (*x*, *z*) + *SHD* (*x*, *z*).

#### Proof

Observe that

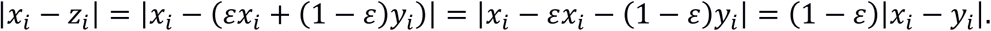

So

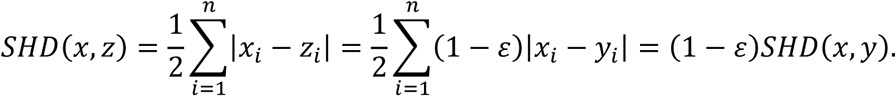

The second equality follows analogously and the third is a trivial consequence of the first two:

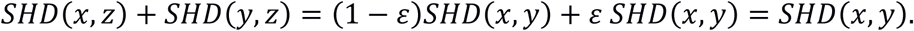

Thus SHD values of admixing populations perfectly represent their common gene pool and admixture ratios.

### MITOMIX: calculation of the best fitting admixture for a given population

Due to the two dimensional surface of earth most populations have several neighbors and at each contact zone admixture may happen between the contacting populations. As we have seen the SHD values between admixing populations are proportional to the admixture ratio of the two populations. Thus we can extend our model to consider an arbitrary number of admixing populations. Let us consider *k* additional populations *y*_1_,…,*y_k_* where 
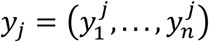
 and 
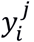
 is the frequency of *i*-*th* mt Hg in the *j*-*th* population.

a. Our first question is whether population ***x*** can be fully obtained from the (linear) combination of other populations, that is whether there are non-negative numbers *ε*_1_,…,*ε_k_* such that *ε*_1_+…+*ε_k_* so that

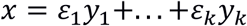

In other words whether *x* can be obtained as the convex combination of *y*_1_,…,*y*_k_ vectors.

a. In theory the above equation can only be solved if all individuals are genotyped from each admixing populations. In practice this is usually not possible, since only a subset of individuals are genotyped and thus leading to sampling bias, moreover often not all admixing participants are represented in the dataset. Thus it is practical to search for the best admixture of *k* populations giving the most similar value to *x*; that is minimizing the SHD value. In particularly we are looking for the linear combination *ε*_1_,…,*ε_k_* where the distance

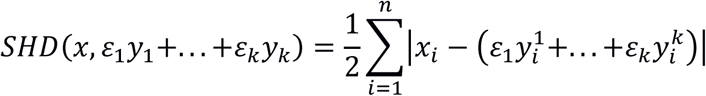

is minimal,

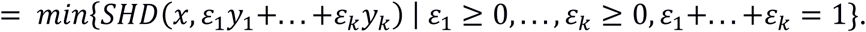

In other words we are looking for those coefficients *ε*_1_,…,*ε_k_* for which the proportional combination of the *y*_1_,…,*y_k_* populations yields the closest SHD value to population *x*.

To compute all possible combinations and proportions of *k* populations in order to find the best fitting admixtures with the smallest SHD values from a test population we implemented a finite software solution. The software is implemented to use multiple threads to speed up calculations on multicore CPU systems. The output file for each analyzed population is a tab separated file that – besides the pairwise best fit population and the corresponding SHD value – also contains an ordered list of SHD values with the corresponding population names and admix percentages, which are integers between the range of 1-100.

### Evolution of the mtDNA tree: haplogroup frequency shift

From an inheritance perspective, the mitochondrial genome can be viewed as a single hypervariable haploid locus, since all of the variants are inherited together (100% linkage) from mother to offspring. As mutation can happen at any mtDNA position and there is no recombination, these aspects hold additional information not available for autosomal traits. From the observed combinations of mutations their consecutive appearance in time can be deducted, which is used to build the mtDNA tree. The mtDNA tree is not static, on a large enough time scale new mutations are fixated giving rise to new “leaves” on the tree, which can expand and spread in the population. When a new mt Hg appears in a noticeable fraction in a given population (gets fixated), by definition the frequency of the other mt Hgs decrease, as the sum of all mt Hg frequencies always equals 1. This phenomenon is detected in time as an mt Hg frequency shift, because part of the individuals of the population will carry the newly fixated mt Hg at the expense of other mt Hgs in the same population (Fig. 1).

**Fig. 1.**
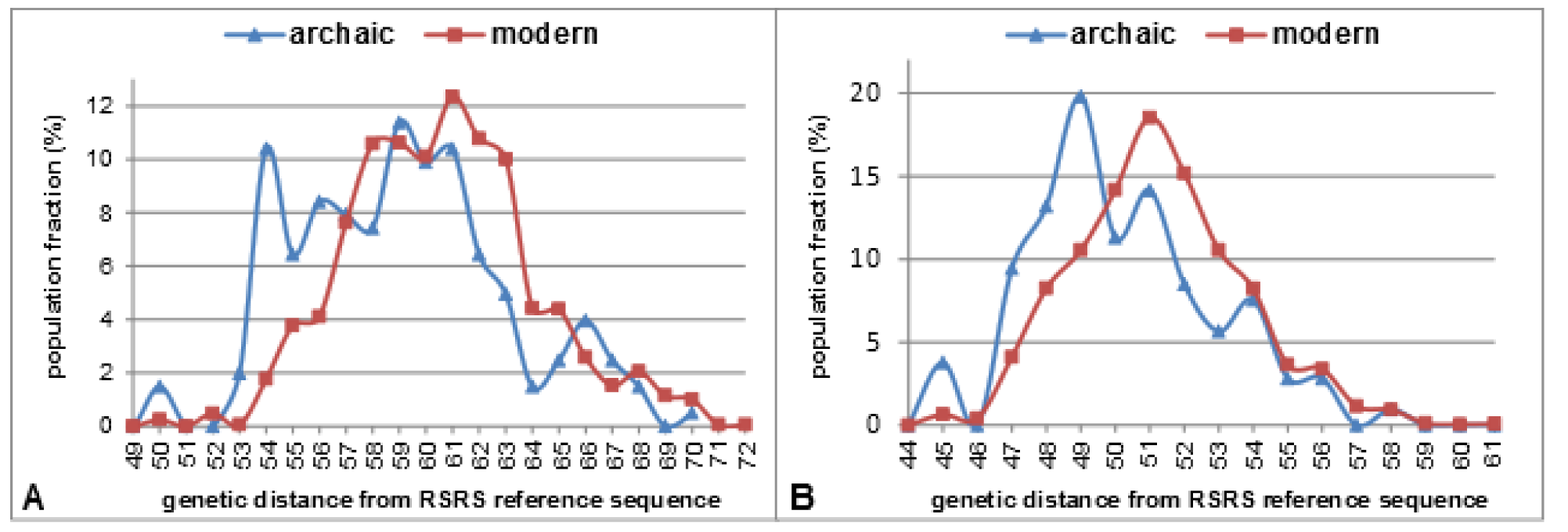
Mitochondrial haplogroup frequency shift between ancient and modern Eurasian populations shown for the most frequent major haplogroups U (**A**) and H (**B**) from available data. Nucleotide distances were calculated from RSRS considering mt Hg defining SNP-s of the current build 17 RSRS mt DNA tree.

As a consequence the frequency of ancestral mt Hgs will stochastically decrease in time. Even though the mutation rate across the mtDNA genome varies, all mutations (except very rare reversion to parent mt Hg) may result in new mt Hgs in case they get fixated. Thus the evolution speed of mtDNA branches are independent of the parent mt Hg itself. Considering constant fixation rate it is expected that the evolutionary speed of different sub-haplogroup branches are proportional to the abundance of their ancestral lineages. For example if a theoretical population contains 99 million people with H, and 1 million with H1 mt Hgs, the chance of a new mutation occurring in H carrying individuals is 99 fold greater than that in H1 individuals. Thus there is 99 fold higher chance that a new H2 than a new H1a haplogroup appears and gets fixated. Although the ultimate mt Hg frequencies are also influenced by population dynamic factors such as population size, expansion, migration and drift the above correlation is well detectable in the distribution of archaic and recent mt Hg frequencies (Fig. 1). One can observe on Fig. 1 that in agreement with our hypothesis the observed frequency shift is less pronounced at rare mt Hgs, located at the extremes of the curve and is most prominent at the most abundant wide spread mt Hgs (middle of the curve).

It cannot be ignored that on longer time scales sequence evolution inevitably alters the mt Hg distribution of populations, which shall bias SHD calculations. The steady decrease and loss of ancient progenitor mt Hgs in time could mask evolutionary distant relations either between ancient and modern or long isolated modern populations, in which case this effect has to be considered. The good correlation between the distribution of archaic and recent mt Hg frequencies indicates that the random population dynamic effects are less pronounced on widespread, older, multicultural mt Hgs which carried by the majority and a standard correction could be applied that reverses the observed distribution shift.

### Reversing Hg frequency shift to correct mtDNA haplogroup frequencies in time

Reversing the mt Hg frequency shift should theoretically result in more accurate estimates of paired distances between modern and archaic populations. Besides, this correction should also improve the distance measurement between modern populations, since it would consider some relation between progeny and progenitor lineages, and thereby revealing evolutionary distant population relations, which are otherwise not detected.

Comparison of available Eurasian recent and ancient mt Hg distributions reveal that the past ~8000 years of sequence evolution has shifted the majority mt Hgs approximately two sub-branches towards the “leaves” on the mtDNA tree (Fig. 1). Considering that at each generation the degree of frequency shift for all mt Hgs is proportional to the frequencies of their parental lineages and the absolute number of people carrying them, an iterative algorithm can be applied to consider this effect, whereby the shift can be reversed. Accordingly, we implemented an iterative method that reverses the mt Hgs frequency shift and calculates the estimated mt Hg frequencies of each population back to an arbitrary date.

Based on the estimated World population sizes [7, 8], we calculated the average population size (P_AVG_) for every generation backwards, using the trapezoid area formula and a constant generation length of 25 years. Based on the current build 17 RSRS mt DNA tree [2] we determined the mutation distances of each mt Hgs from their parental Hgs. In cases where this distance was more than one nucleotide, we added intermediate parental Hg(s), hence all mt Hgs were rendered to one mutation distance from their parental lineages. Then from our newly assembled database containing 12261 individual samples from 62 modern populations and 629 samples from 25 archaic populations we calculated the fraction of Hg-s with identical mutational distances from RSRS, thereby we defined (*k*) distance layers on the mtDNA tree.

We denoted the absolute mutational distance of an mt Hg from the RSRS reference sequence with *d_RSRS_* (*Hg*) and the allele frequency of the mt Hg with *AF*(*Hg*). The overall fraction of people carrying mt Hgs at a given mtDNA distance layer (*OVERALLFRACT*_*k*_) corresponding to a certain time interval can be expressed as:

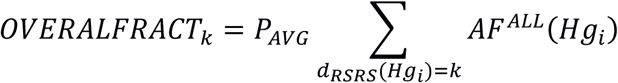

Considering random/constant mutation and fixation of mt Hgs the amount of new leaves are directly related to the abundance of their parent mt Hgs. Thus we computed the parental mt Hg fraction of each mt Hgs for each population 
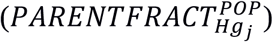
 from the estimated number of people at a given date carrying mt Hgs at the same RSRS distance (*OVERALLFRACT*_*k*_
) and the allele frequency of the given mt Hg in the population by the following formula:

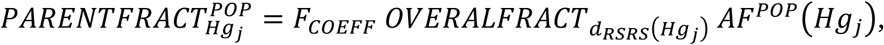

where F_COEFF_ is an experimental value, representing a constant fixation coefficient. Starting from present day, the dataset can be iterated step-by-step for each generation backwards using the above formulas and for each population originating from newer or spanning dates with the iteration step’s date. As a result, at each iteration step a fraction of ancestral mt Hgs is added to the mt Hg pool of the populations at the expense of their progeny lineages.

### mt Hg frequency shift correction of populations

To assess the time dependence of mt Hg distribution shift a representative number of mt Hgs are required that originate from similarly dated archaic populations. Since the number of available archaic mitogenomes is restricted, we attempted to balance between the number and age similarity of archaic samples to achieve these criteria. Therefore for this calculation we excluded Paleolithic, Armenian Iron Age and Hungarian Conqueror samples from the dataset, as their population age largely deviated from other archaic populations. Using an iterative approach we transformed the mt Hg distribution of our modern populations and reversed the mt Hg shift to the average population age of our selected archaic populations back to 8000 years ago, and the corrected mt Hg distributions were used throughout the experiments. Experimentally we determined the best fitting coefficient (F_COEFF_=0.000125) which corrected modern mt Hg distribution to best match the mt Hg distribution of available archaic populations (Fig. 2).

**Fig. 2.**
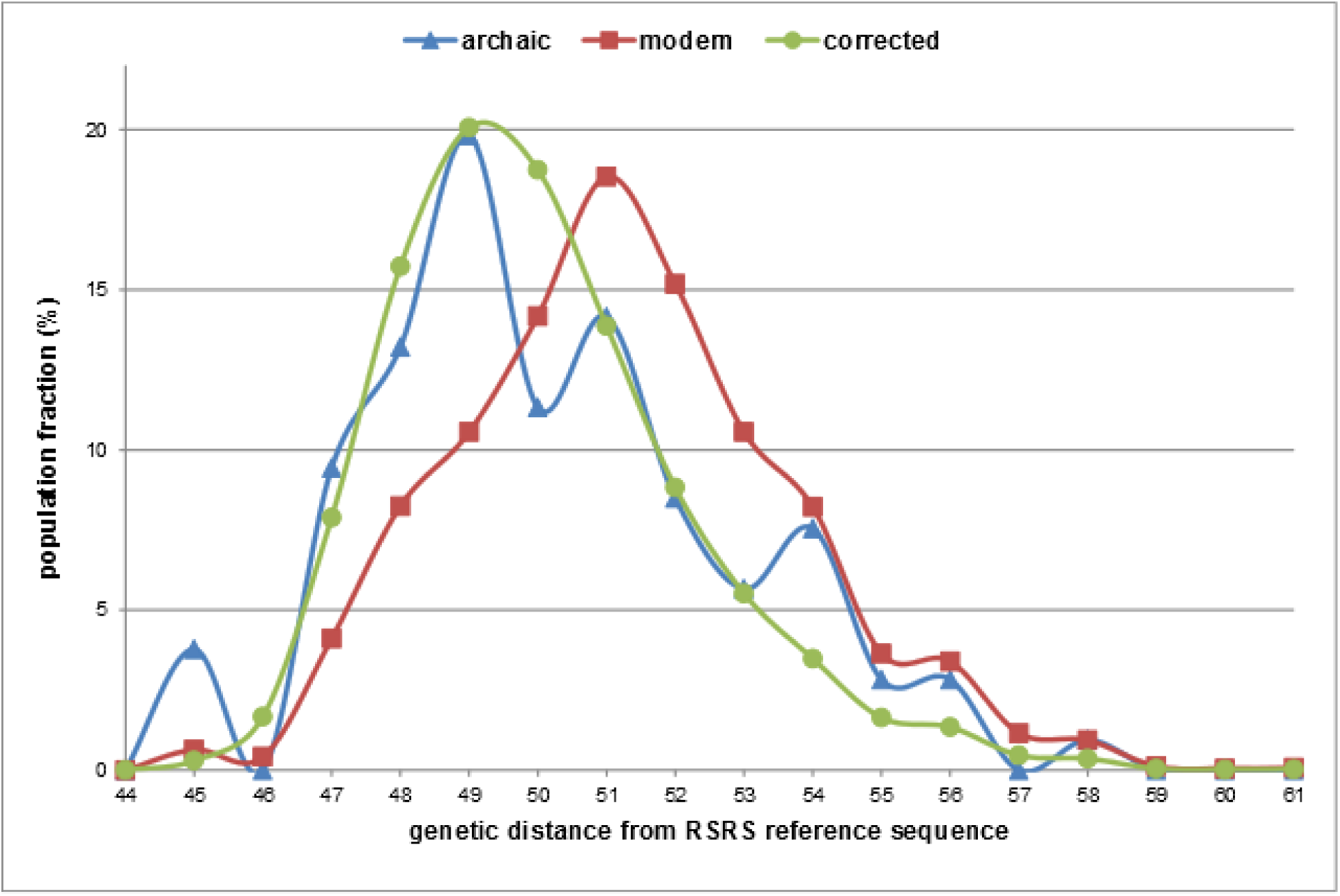
mt Hg frequency shift correction of modern major mitochondrial haplogroup H back to 8000 years ago, using best fitting coefficient (F_COEFF_= 0.000125).

To illustrate the effect of the mt Hg shift correction at the level of individual populations we compared the original and corrected mt Hg frequency data (Supplementary Table 2) in a subset of selected modern and archaic populations. We visualized the age distribution of the original and corrected mt Hgs (Fig. 3).

**Fig. 3.**
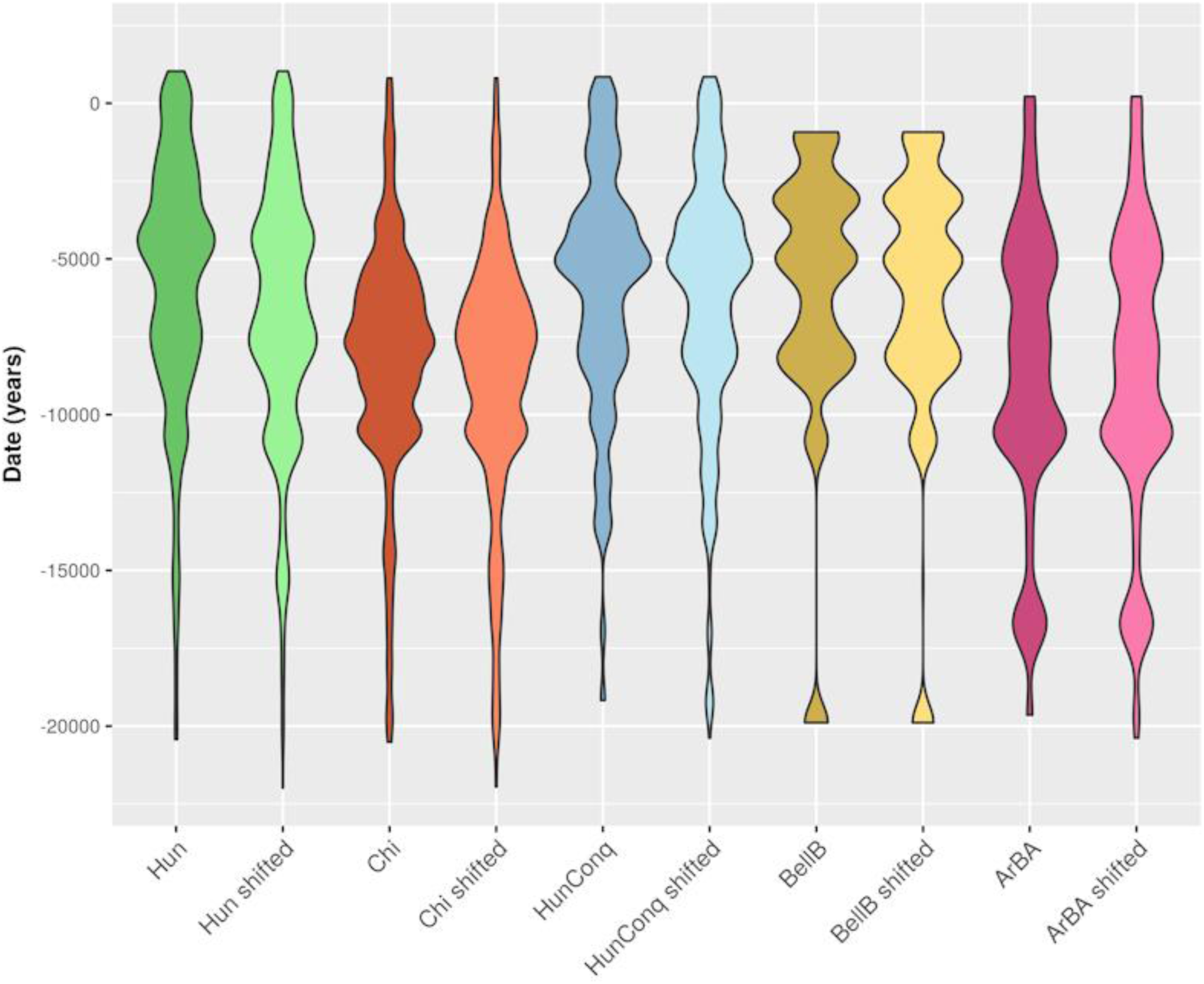
mt Hg age (after Behar) distribution of a selected modern (Hun – Hungarian, Chi – Chinese) and archaic (HunConq – Hungarian Conqueror, BellB – Bell Beaker, ArBA – Armenian Bronze Age) populations before and after mt Hg shift correction.

Fig. 3 visualizes that in accordance with our assumption each population has a unique distribution of mt Hgs. Fig. 3 also demonstrates that the applied correction does not alter the original distributions significantly and populations remain most similar to themselves after the correction. The correction is most pronounced at abundant mt Hgs while it is negligible at infrequent mt Hgs corresponding to the observations of experimental data. Furthermore, the effect of correction is less pronounced on archaic populations, because being older they were iterated for fewer generations, and also because corresponding to the World population estimates, the average population size used in the iteration formula was smaller.

### Effect of sample size on SHD calculations

The exact SHD value can only be calculated from complete population data; however in practice from most populations just more or less representative numbers of samples are available. Thus, the calculated SHD value is inevitably distorted by sampling bias, the extent of which depends on the number of samples and the mt Hg diversity of the population. We calculated the inverse Simpson index 
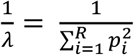
[9] representing the mt Hg diversity for each studied population (Supplementary Table 3). The mt Hg diversity values scatter in an extensive range among the analyzed populations. It is notable that populations with the lowest mt Hg diversities are usually isolated by natural and/or cultural barriers and are often small in number: many of the small North-East Asian populations, the Basques or populations situated on islands such as Japanese and Sardinians belong to this group. An extreme example is the Commander Island population (Com) consisting of only two mt Hgs. Among the most diverse populations in our database are the Chinese (Chi, a group with many subpopulations) and many of the European populations, especially the ones with complex ethnogenesis and/or those surrounded by a number of neighbors with diverse genetic origin. The archaic samples show generally smaller diversity that is not only the result of low sample size, but also derives from the recent exponential expansion of the world population, having caused an unprecedented burst in mtDNA evolution.

Theoretically representative sampling of populations with high Hg diversity –where mt Hg count is comparable to sample count– requires a larger sample set than that of less diverse populations. In order to assess the effect of sample size on SHD calculation, we performed a Monte Carlo simulation [10] on experimental data. For this analysis a highly diverse (Italian), a moderately diverse (Danish) and a less diverse (Japanese) population had been selected from our compiled population database. Two subsets of samples (without replacement) were randomized 1000 times from each population and the average SHD values between the two subsets were calculated for different sample sizes (Fig. 4).

**Fig. 4.**
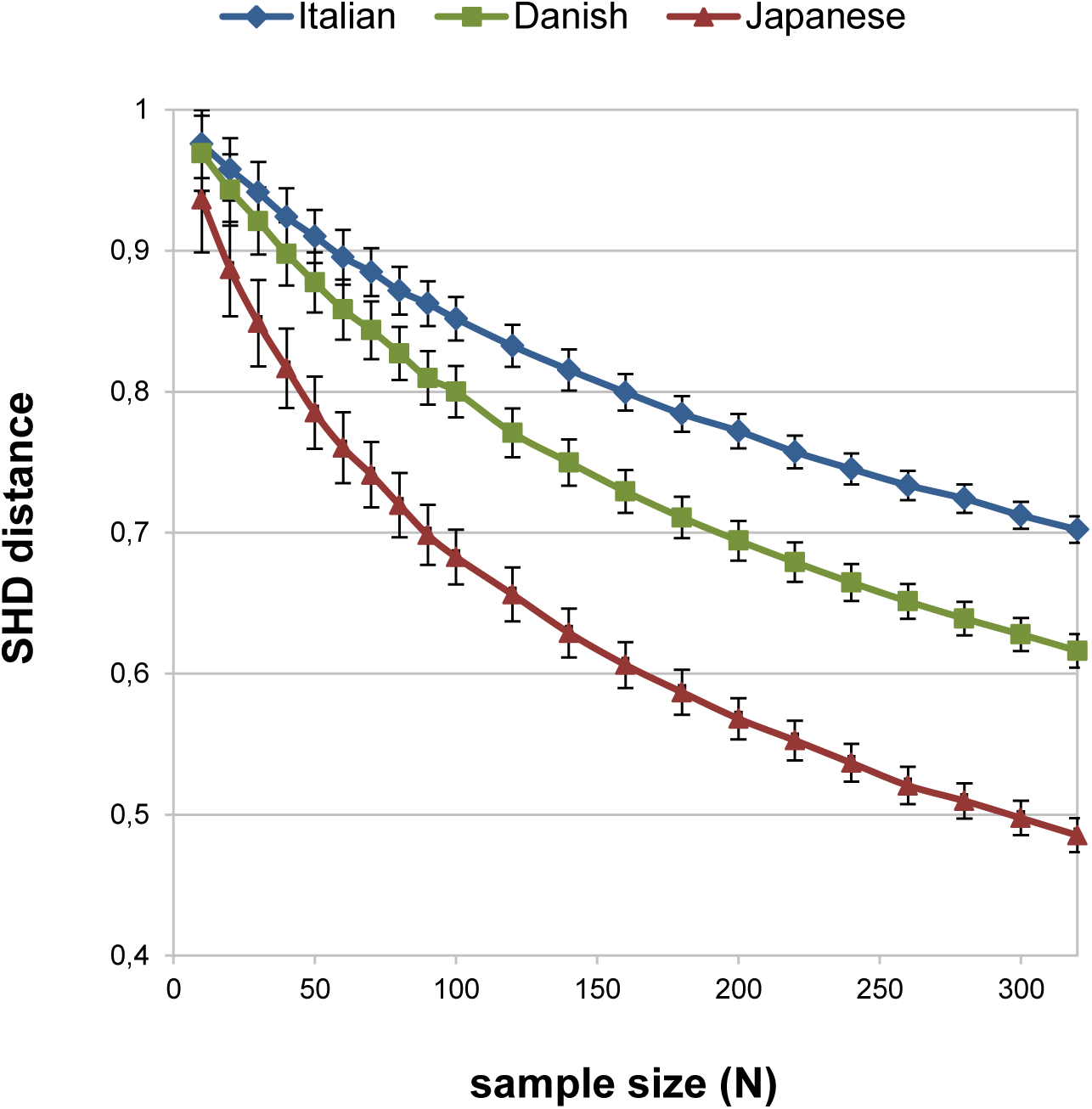
The dependence of calculated SHD value on intra population diversity and sample size. The error bars represent the standard error of the calculated SHD values of 1000 random iterations.

Our analysis demonstrates that in case of diverse populations even high sample size results in relatively high SHD values, e.g. considering two random Danish cohorts (n=300) gives 0.62 average distance from each other (in case of highly diverse Italian the calculated distance is 0.70 while between the less diverse Japanese is 0.48), instead of the theoretical zero. This raises the question as to what extent between populations values are distorted, and how much are they informative? To answer this question, next we performed a similar Monte Carlo simulation with populations of various diversity and mt Hg overlap. We analyzed two populations without significant shared mt Hgs (Danish, Japanese), two populations with somewhat similar mt Hg distribution (Italian, Danish), and two populations with similar mt Hg distribution (Danish, Hungarian) by calculating the SHD values between random subsets (sampling without replacement) of the above populations with different sample sizes (Fig. 5).

**Fig. 5.**
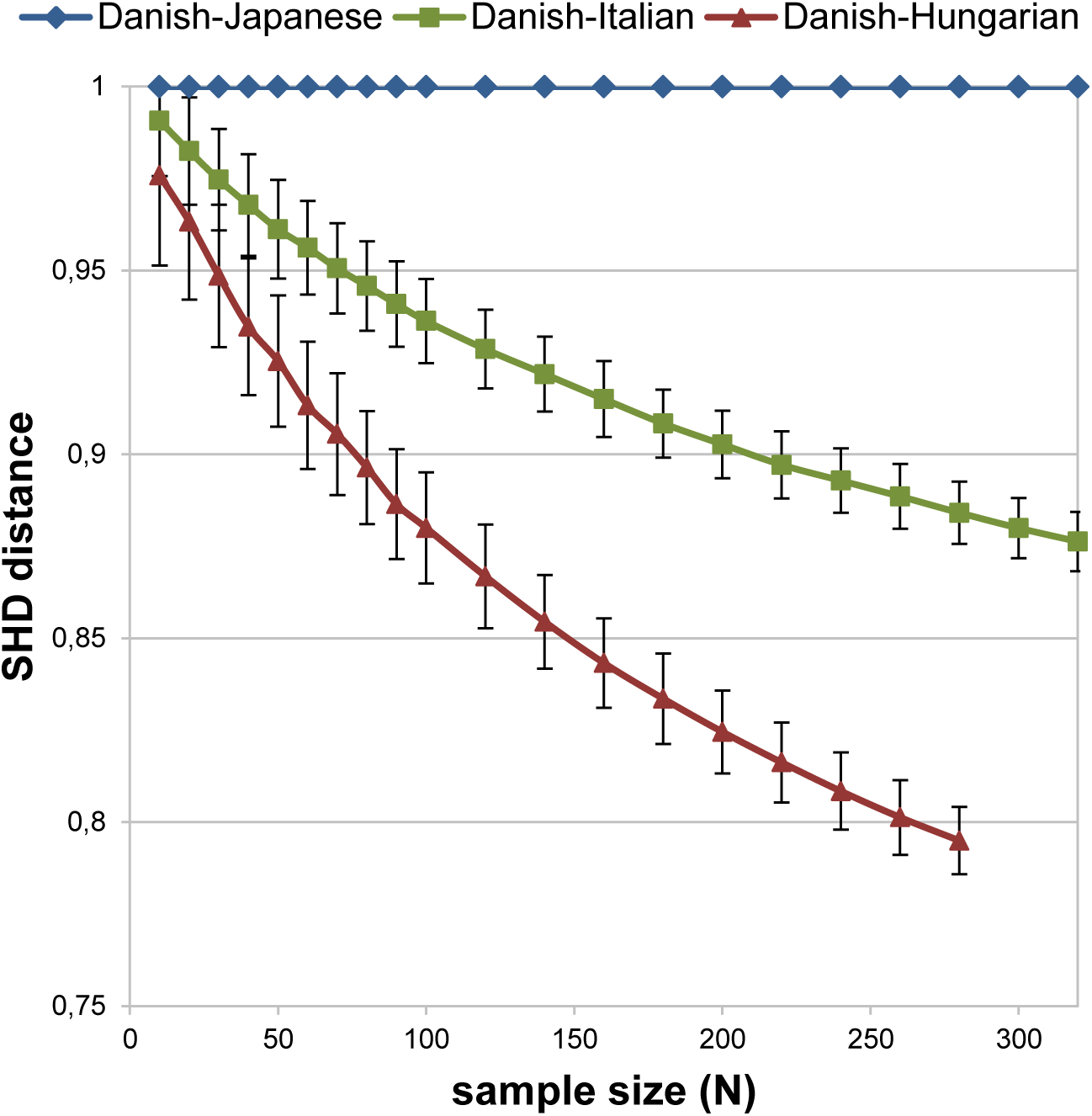
Dependence of the SHD value on inter-population diversity and sample size. The error bars represent the standard error of the calculated SHD values of 1000 random iterations.

The result of the simulation indicates that distortion of SHD values are indeed proportional to sample size, but in spite of this bias the *relative* SHD values between population pairs are statistically well represented, meaning that the distance order between the 3 populations (Danish –Hungarian<Italian<Japanese) remains the same at any sample size. This result is due to the fact that random sampling statistically does not disturb the ratio of shared and non-shared mt Hgs. Consequently, the calculated SHD value will be lowest between populations with most similar mt Hg distributions and the order of SHD values will statistically correspond to the same relative order of similarity regardless of population diversity, while as expected, the standard error depends on the number of samples and the fraction of shared mt Hgs.

Furthermore, comparing low and high mt Hg diversity populations the bias and the correct number of samples required for the analysis mostly depends on the representativeness of the less diverse population, since relatively low sample size from the more diverse population is expected to result mostly a large number of non-detected but non-shared Hg-s, which does not influence the calculated distance value. We can also infer from Fig.s 3 and 4 that in case of diverse populations relatively high SHD values like 0.8 in case of 100 samples or 0.6 in case of 300 samples, can be interpreted as very high levels of similarity. The correlation between SHD values with sample size and population diversity raises the possibility of a theoretical correction by Monte Carlo simulation to estimate a more realistic SHD value from available data. However we caution that such a correction will inevitably “overcorrect” distances with very low sample sizes because in this case the assumption that the sample subset represents the whole population’s mt Hg diversity and distribution is obviously not true, thus it will distort the *relative* order of SHD values – a very valuable attribute of the method - in an unpredictable manner. For this reason we did not correct our data, but we present an estimation of expected average SHD values at the given sample size for each of the populations based on the assumption that their observed mt Hg diversity and distribution is representative (Supplementary Table 3). We also implemented this Monte Carlo simulation method in the software package.

### SHD and F_ST_ calculations from experimental data

Using our mitogenome database assembled from available data we calculated the SHD matrix of all modern and archaic Eurasian populations (Supplementary Table 4), and then from this distance matrix, we created a hierarchic cluster of the analyzed populations (Fig. 6).

**Fig. 6.**
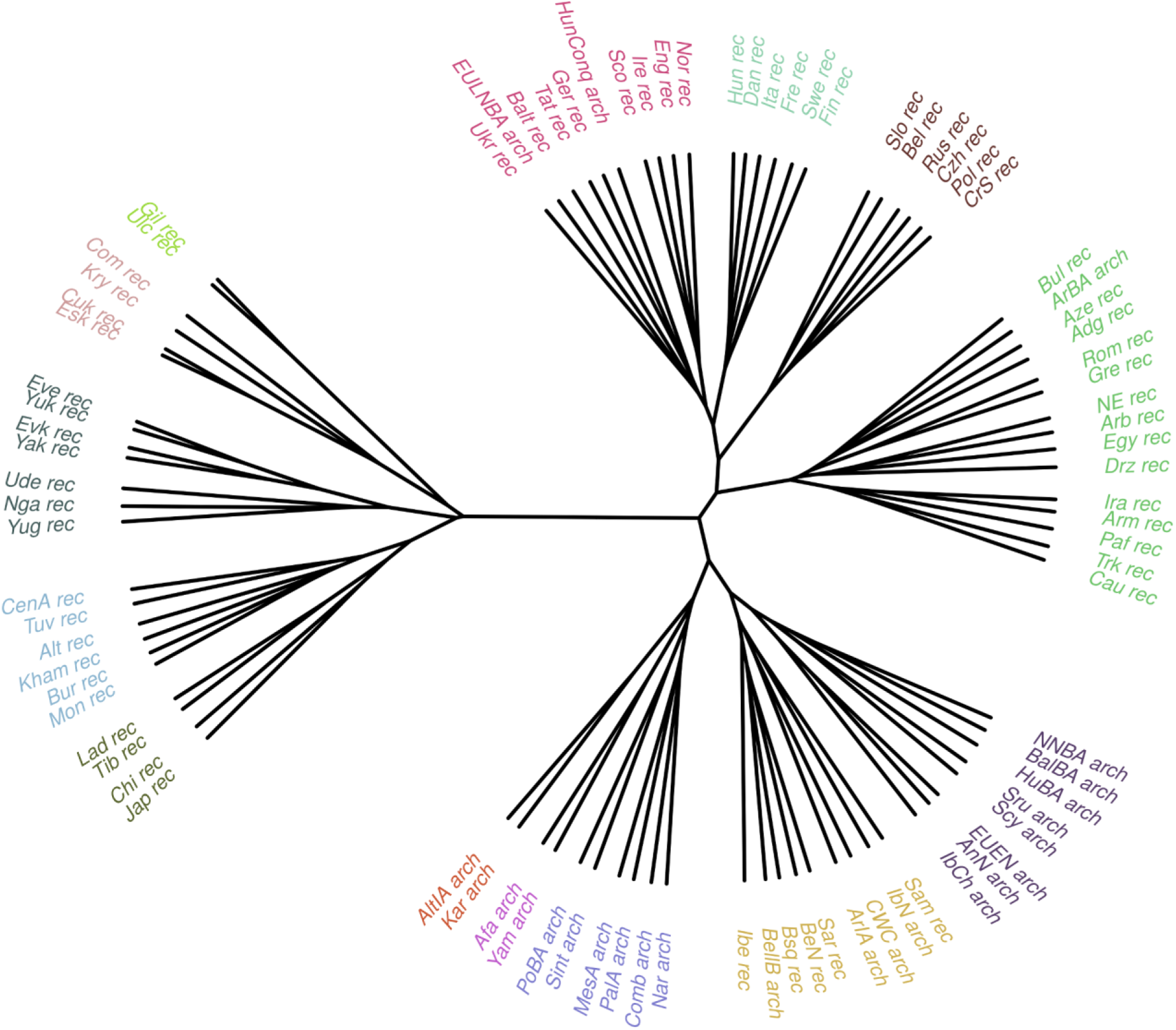
Unrooted hierarchic cluster of modern and archaic populations based on the SHD matrix.

The structure of the tree seemingly sequesters genetically similar populations into the same branches, and separates modern populations mostly according to their geographical locations. The cluster contains 2 main sub-clusters; Asian populations on the left, European populations on the right section. Within the European group most ancient populations are sequestered into a large sub-cluster (center bottom section in Fig. 6) while modern populations are separated mostly according to their geographical locations.

The Asian group is also separated into 5 sub-clusters based on their close geo-localizations, in agreement with previous data [11]. These are **1)** the east coast of Russia, and just north of Japan (Gilyak and Ulchi populations light green); **2)** Northeastern Siberia (Commander Islands and Koryak, Chukchi and the Eskimo /Naukan + Aleut + Inuit+ Tlingit/ populations in pink); **3)** Siberian/Russian Far East (Even, Yukaghir, Evenki, Yakut, Udegei, Nganasan, Mansi, Khanty in dark blue); **4)** Central Asia (composing Central Asians /Uzbek + Kyrgyz + Kazakh + Turkmen/, Tuvan, Altai region /Teleut + Shor + Kizhi + Tubular/, Khamnigan, Buryat, and Mongol in light blue); **5)** and East Asia (Ladakh, Tibetan, Chinese, Japanese in black. The SHD based unrooted hierarchic cluster even depicts the complex genetic relationship among neighboring Siberian populations, which indicates assimilation between North Tungusic (Evenk and Evens) and Amur Tungusic (Yukhagir, Yakut) populations as has been shown previously [12].

The modern European group is divided into 4 main clusters: **1)** modern Slavic populations, known to have common genetic origin [13] (Slovakian, Belarus, Russian Czech, Polish, Croatian/Serbians dark brown); **2)** Finnish, Swedish, French, Italian, Danish and Hungarian populations in light blue; **3)** Norvegian, English, Irish, Scottish, Hungarian Conquerors, German, Tatars, Baltic /Lithuanian + Latvian + Estonian/ populations, Central_LNBA and Ukraine in dark magenta. **4)** the last cluster consists of 3 further subgroups (in green) including **4a)** Trans-Caucasus region (/Chechen + Kurdish + Karachay + Georgian + Ossetian + Kabardinian + Cherkess + Darginian + Dstan + Kalmyk + Avar/, Turkish, Pakistan + Afghanistan, Armenian, Iranian populations); **4b)** Middle East (Iraqi + Lebanese + Jordan + Palestine/, Saudi Arabia, Druze, Egyptian populations); and **4c)** the Black-Sea Region (Greek, Romanian, Azeri, Adgey, Bulgarian population, Armenian-Iran Chalcolithic-Bronze Age). All these data indicate complex relations between the European populations, which is poorly represented on a tree, as the genetic distances alone do not enable uncovering the fine details of complex admixtures.

In spite of the limited data available from ancient populations the tree corresponds well with other known genetic data, due to the reliability of relative SHD values as discussed above. This is best demonstrated by the Afanasevo Yamnaya affiliation, since just 5 Afanasevo and 15 Yamnaya mitogenomes were available, nevertheless the tree reflect their known close relations [14]. In general most ancient populations fall into expected branches, and are more similar to each other than to modern populations, as they contain mainly ancient Hg lineages, while most modern populations are full of terminal sub-groups due to recent population expansions. However, some modern populations are clustered together with archaic cultures, most notably Basques, Sardinians, Saami and Iberians, many of which had formerly been identified as “genetic outliers” carrying larger portion of ancient genetic elements [15, 16]. This sub-branch seems to support relations between Iberian Neolith, Bell Beaker, Corded Ware cultures and the modern Iberians, Basques and Sardinians as suggested before [17].

Seemingly the tree shows only a few incorrect classifications, which can be explained by sampling bias, insufficient sample numbers from diverse populations. One example is Germans (Ger) which were ordered closer to Volga Tatars (Tat) than to their European neighbors (Fig. 3), as presumably the 105 German samples do not properly represent this genetically diverse group.

In order to compare we also calculated the F_ST_ population distance matrix (Supplementary Table 4) and performed the corresponding hierarchic clustering (Supplementary Fig. 1) for the same populations (except Baltic Late Bronze Age (BalBA) for which fasta files were not yet available). Comparing the hierarchic clusters of SHD and F_ST_ data it is apparent that the major topology of the two clusters shows great extent of similarity. Besides the similarities there are also notable differences between the two clusters, for example SHD incorrectly distorts evolutionary distances, and curtails branch lengths between European and Asian groups. This is an expected consequence, since by definition the distance is 1 between any two populations with no shared mt Hgs regardless of the mutational distances. There are also notable differences in population groupings; one typical example is the classification of Polish people which are placed to a common sub-branch with Finnish by F_ST_, while SHD places them into a common sub-branch with all other Slavic populations (Russian, Serbian, Croatian, Czechs, Slovakian and Belarusian). The latter is a more realistic grouping, as according to historic and genetic data, the Polish people genetically belong to Slavs [18]. Examining the detailed relation of Polish to other populations it appears that 29 non Slavic populations fall within the Polish – any other Slavic population F_ST_ distance range (0.015-0.042), 7 of which are even closer to Polish than the closest Slavs, Croatians/Serbians. In contrast, the closest fits to the Polish population with SHD (Supplementary Table 4) are Croatians/Serbians and Russians while only 7 non Slavic populations fall within the SHD range of Polish - other Slavs (0.754-0.876) and all of these are represented among the 29 detected by F_ST_. Nevertheless, distance order of Slavs is just the same with both methods. This example well illustrates that at smaller genetic distance ranges F_ST_ calculations give just tentative results while SHD based calculation is more consequent and provides better resolution.

Another good example is the clearly admixed Hungarian Conqueror population, in which SHD distinctly indicated an East Eurasian component, which was not detected by F_ST_, and instead positioned the Conqueror population between European and Asian groups. In contrast when the Asian subset of the Conquerors was treated as separated subgroup, both methods detected a similar set of related groups. This example well demonstrates that SHD is superior for detecting admixtures, which is very relevant information for tracing population histories.

### MITOMIX analysis of experimental data

Using our database we performed the MITOMIX analysis of 25 archaic Eurasian and 62 modern populations (Supplementary Table 5) without any preselection of potentially admixing partners. MITOMIX generally indicated admixture between geographically neighboring populations which are frequently grouped together by the hierarchical clustering method using SHD. Geographical co-localization of the test populations with populations contributing to the hypothetical admixtures clearly indicates that local admixture has been a predominant process throughout human history. A good example for this are the Evenki – Even – Yakut – Yukhagir [19–21] or the Eskimo – Chukchi - Commander Island admixtures [22] corroborating previous genetic data. The analysis of Commander Island population also demonstrates that MITOMIX do not force distantly related populations into the best hypothetical admix which do not improve the fit of the model. In this case the best admix indicated 100% Eskimo population as the source, suggesting a genetic drift [23].

In addition the MITOMIX algorithm also correctly identifies relations between populations that are classified into different sub-branches by the hierarchic clustering of SHD matrix. One good example is the Hungarian population, which is genetically very different from the surrounding Slavic groups and accordingly is clustered on different branches (Fig. 6); however MITOMIX consequently identifies varying level admixture components of modern Hungarians from their neighboring Serbian/Croatian, Slovakian and Romanian populations, which are known to have a common history with Hungarians. A few other examples of correctly identified admixtures by MITOMIX are the indication of Finnish-Sami [24]; Italian-Sardinian [25, 26] or Italian-Armenian admixture components that had been detected previously at the genomic level as well [27].

Our dataset contains SHD and MITOMIX data for all 87 populations (Supplementary Table 4-5), but below we discuss in detail only the population genetic analysis of the modern Hungarians.

### FST, SHD and MITOMIX analysis of modern Hungarians

The SHD matrix of the available 87 populations shows that modern Hungarians have the closest SHD values to the modern Danish (Dan 0.684), Italian (Ita 0.741), Dutch/Belgian (BeN 0.743), Basque (Bsq 0.771) Polish (Pol 0.795) and Finnish (Fin 0.798) populations (Supplementary Table 4), and the first three of these are also the closest according to F_ST_ distance values (BeN 0.00175, Dan 0.01095, Ita 0.01203) (Supplementary Table 4). Hierarchic clustering of the SHD matrix (Fig. 6) positions modern Hungarians into a common European subgroup with modern Danish, Italian, French, Swedish and Finnish populations. The closest ancient populations to modern Hungarians by SHD are Baltic Late Bronze Age (BalBA 0.815), Bell Baker (BellB 0.830) and European Early Neolithic (EUEN 0.879).

However, our MITOMIX analysis (Supplementary Table 5) reveals that the shared mt Hg distribution of modern Hungarians are most similar to a hypothetic mixture of Dutch/Belgian (BeN 32%), Danish (Dan 22%), Basque (Bsq 17%), Serbian/Croatian (CrS 11%), Baltic Late Bronze Age archaic (BalBA 8%), Bell Beaker archaic (BellB 5%) and Slovakian (Slo 5%) populations resulting in 0.6015 SHD value. Alternative, similar probability MITOMIX combinations (indicating of the same main contributors) also include low contributions of German (Ger 4-6%), Northern Neolith Bronze Age (NNBA 3-4%), Hungarian Conqueror (HunConq 3%), Srubnaya archaic (Sru 3%), Romanian (Rom 2-3%), and Yamnaya archaic (Yam 1-2%) populations. As modern Hungarians are one of the most diverse populations (Supplementary Table 3) Monte Carlo simulation predicts that from the analyzed modern Hunagarian samples SHD value 0.48 would be close to the minimum (Supplementary Table 3), thus the empirical 0.6015 SHD value from the theoretical admixture indicates that it accounts for the origin of the majority of mt Hg components in modern Hungarians. Neighboring nations together contribute around 16% to the best hypothetical admix; while the largest admix components surprisingly are derived from geographically distant modern European populations.

It is important to note that mt Hg sharing can only stem from common ancestors or admixture, and therefore these data indicate that modern Hungarians either must be descendants of ancient populations that have experienced significant admixture with ancestors of modern Danish, Dutch/Belgian, Italian and Basque people or have co-ancestry with the aforementioned populations. As the occurrence of such admixture can be excluded in modern history it must have taken place in prehistoric times, and if so, mt Hg overlap with these populations should be confined mostly to older mt Hg lineages. To test this hypothesis we assembled a detailed list of mt Hgs from the best MITOMIX (Supplementary Table 6). Based on this data, we created the density plot of the estimated age of shared mt Hgs for the Hungarians (Hun), the best hypothetical admix (mixFreq) and the populations contributing to this admix (Fig. 7).

**Fig. 7.**
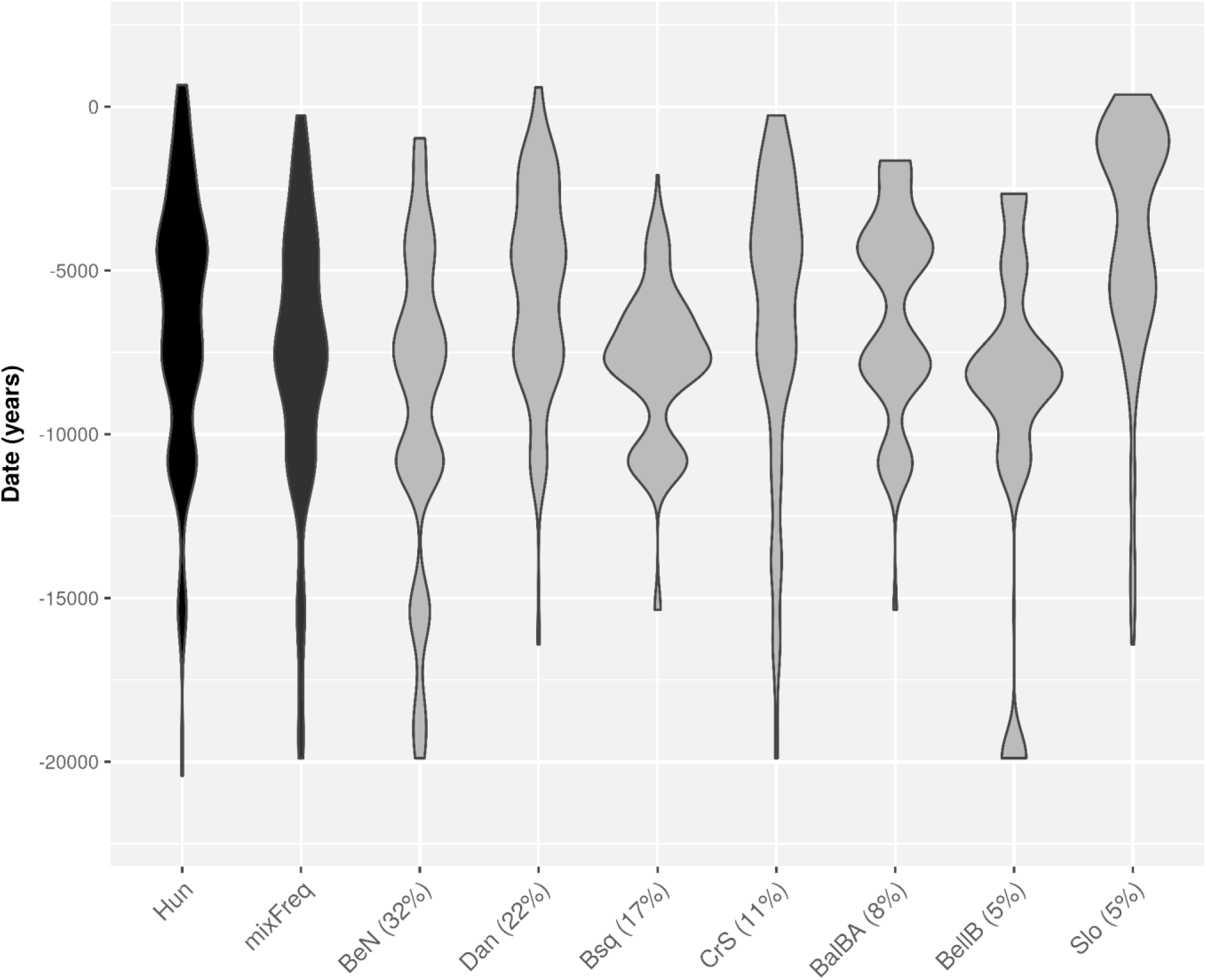
The estimated age distribution of the shared mt Hgs between Hungarians (Hun), the best hypothetical admix (mixFreq) and the populations contributing to this admix: Belgian/Dutch (BeN), Danish (Dan), Basque (Bsq), Croatian/Serbian (CrS), Baltic Late Bronze Age culture (BalBA), Bell Beaker culture (BellB), Slovakian (Slo). The numbers in parentheses indicate the contributions to the best hypothetical admix.

The hypothetical admix population has a remarkably similar mt Hg distribution compared to the Hungarians affirming the efficacy of the MITOMIX algorithm. The estimated age distribution of shared mt Hgs shows that main shared components with Basque and Bell Beaker Cultures are the oldest, mostly originating from the Paleolithic/Neolithic period. Shared mt Hgs with the Baltic Late Bronze Age culture and Belgian/Dutch populations mostly originate somewhat later, around the Late Neolithic and Bronze Age. In contrast, shared mt Hgs with Serbians/Croatians, Danish and especially Slovakians have a wide age distribution including young mt Hgs originating from the Iron Age or later. The average mutational distances (measured from the RSRS reference sequence) of the mt Hgs shared between Hungarians and these populations were the following: Basque 48.6; Bell Beaker, 48.7; Baltic Late Bronze Age 49.5; Belgian/Dutch 50.8; Serbian/Croatian 51.3; Danish 52.7; Slovakian 55.4. These values also show that the average RSRS distance of shared Hgs between Basque, Belgian/Dutch and modern Hungarians are comparable to that of Bell Beaker and Baltic Late Bronze Age cultures, which indicates that they indeed originate from an ancient European source, while Serbian/Croatian, Danish and Slovakian admixture are probably more recent. The remaining mt Hg fraction (~22.8%) of modern Hungarians that were not found in the best admix combination is summarized in Supplementary Table 6, ordered by the fraction of mt Hgs shared with Hungarians. There are 59 populations with shared mt Hg fraction ranging between 0.04-6.49 percent. Notably Italians, Armenians, Iranians, and Pakistanis + Afghanis have over 4 percent shared mt Hgs with modern Hungarians; however their non-shared Hg-s are different to such a great extent that renders them an inadequate fit in the best admixtures for MITOMIX.

To find out the potential common European ancestral sources of the Hungarian, Basque, Belgian/Dutch and Danish populations we also performed a MITOMIX analysis in which only ancient populations were considered as source (Table 1).

**Table 1.** Archaic components of modern populations contributing to hypothetical Hungarian admixture. MITOMIX analysis of Hungarian and other non-neighbor modern test populations (indicated by unrestricted MITOMIX analysis), restricted to consider (archaic) populations dated from earlier or spanning time period only as the source of admixture.

|  | **best fitting admixture of archaic populations (k=4)** |  |  |  |  |
| --- | --- | --- | --- | --- | --- |
| **test population** | **SHD** | **POP 1 (%)** | **POP 2 (%)** | **POP 3 (%)** | **POP 4 (%)** |
| Hungarian | 0.744 | BellB (31) | BalBA (31) | EULNBA (21) | AnN (17) |
| Basque | 0.770 | BellB (78) | IbCh (13) | BalBA (9) |  |
| Belgian/Dutch | 0.753 | BalBA (48) | BellB (29) | CWC (12) | IbCh (11) |
| Danish | 0.825 | BalBA (34) | EULNBA (34) | EUEN (21) | IbN (11) |

Out of the 4 major archaic cultures contributing to the best Hungarian ancient MITOMIX three are shared with the other European populations, notably Bell Beaker + Baltic Late Bronze Age with Basque and Belgian/Dutch while Baltic Late Bronze Age/European Late Neolith/Bronze Age with Danish population. Accordingly the closest modern populations to Bell Baker are Basques (0.793), Hungarians (0.846) and Belgian/Dutch (0.849); to Baltic Late Bronze Age are Belgian/Dutch (0.830), Hungarians (0.851) and Basques (0.882); to EULNBA Belgian/Dutch (0.899) has significant affinity, while Hungarians (0.906831) and Danish (0.925) also have some similarity.

It is notable that Baltic Late Bronze Age and Bell Beaker cultures also show up in the Hungarian MITOMIX when both modern and ancient populations are considered together, indicating that these ancient cultures played a prominent role in the formation of the modern Hungarian gene pool. Nevertheless other ancient admixture sources may be missing from the dataset as ancient admixtures result in significantly higher (0.744) SHD value compared with the unrestricted admixture (0.602).

The above data suggest that a significant fraction of the modern maternal Hungarian gene pool may have originated as old as from Neolithic-Bronze Age, in which case those old mt Hgs are expected to persist in the modern population. Although our ancient database contains just 10 samples from Bronze Age Hungary (Hungarian Bronze Age, HuBA) we tested the co-occurrence of these 10 mt Hgs in other archaic and modern populations (Table 2).

**Table 2.** Co-occurrence of Hungarian Bronze Age mt Hgs. Distribution of mt Hgs found in Hungarian Bronze Age archaic samples in the analyzed populations. The fixation dates are based on Behar et al [6].

| **Hungarian**<br>**Bronze Age**<br>**mt Hg** | **fixation date**<br>**Year (SD)** | **mt Hg fraction in other populations** |  |  |  |  |  |  |  |
| --- | --- | --- | --- | --- | --- | --- | --- | --- | --- |
| H11a2 | 5406 (1873) | Hun (1/289) | Dan (1/2834) | Ita (1/645) | Fin (1/651) | Pol (1/300) | Rom (1/31) | Adg (1/24) |  |
| H2a1 | 7713 (1754) | Hun (3/289) | Dan (8/2834) | Ita (1/645) | Bsq (4/453) | Fin (5/651) | Czh (1/64) | Sco (2/60) | Ira (2/394) |
| J1c9 | 4001 (3768) | Dan (3/2834) | Ita (2/645) |  |  |  |  |  |  |
| K1a1a | 9093 (3539) | Dan (2/2834) | Pol (1/300) | Sco (1/60) |  |  |  |  |  |
| K1a2a | 9165 (3986) | Hun (1/289) | Dan (1/2834) | Ita (1/645) | Arm (1/422) | NNBA (1/19) | IbCh (1/21) | IbN (1/11) |  |
| K1c1 | 5215 (1953) | Dan (6/2834) | Fin (7/651) | Sco (1/60) | Eng (1/153) | HunConq (2/102) |  |  |  |
| T1 | 16309 (6538) | Rus (1/233) | Arm (1/422) | Aze (1/65) | Tuv (1/26) | AnN (1/62) | ArBA (1/61) |  |  |
| T2b | 10069 (1610) | Hun (4/289) | **multicultural, found in 22 modern and 8 archaic populations (102/12890)** |  |  |  |  |  |  |
| T2b2 | 6462 (3515) | Hun (2/289) | Dan (2/2834) | Swe (1/107) | Ire (1/92) | Fre (1/141) | EULNBA (1/44) |  |  |
| T2b3 | 9202 (1961) | Dan (1/2834) | Rus (1/233) | Aze (1/65) |  |  |  |  |  |
| U5a2a | 12965 (5917) | Dan (1/2834) | Ita (1/645) |  |  |  |  |  |  |
| U5a2b | 11437 (3186) | Dan (2/2834) | Pol (2/300) | Rus (2/233) | Ukr (1/33) | CrS (1/122) | Bul (1/46) | Bur (2/133) | Scy (1/22) |

Table 2 shows that 5 out of 10 mt Hgs present in Hungarian Bronze Age samples are also found in modern Hungarians. Moreover the majority of these mt Hgs are also found in those populations (Danish, Italian) that were indicated as closest relatives of modern Hungarians by our SHD data, hierarchical clustering and MITOMIX analysis. This finding supports that a significant fraction of mt Hg overlap can be explained by admixture of ancient European populations before the Bronze Age, and that genetic continuity exists between these ancient European sources and modern Hungarians.

## Discussion

The most commonly used descriptive statistics in population and evolutionary genetics are fixation index (F_ST_) and related (G_ST_, Q_ST_) statistics [28–32]. These various formulas measure population differentiation (subdivision) based on the variance of diversity originating from divergence [33]. Because of this theoretical premise, these approaches are best suited to calculate the genetic distance between populations where the effect of divergence is determining. Moreover these methods are based on unrealistic, idealized models of population structure, migration and evolution [34], making some of the inferences drawn from real populations questionable, thus many of the F_ST_ population distance calculations that assume isolated, stationary or equal sized populations [29–31, 35], can lead to bias when these assumptions are not met. Available genetic data from ancient and modern populations clearly indicates that admixture has played a major role in shaping human population structure from as early as the Neolithic [27] period. Therefore we argue that in case of closely related and/or admixing populations SHD based population distance calculations give more realistic results and provide better resolution than F statistics (as demonstrated on the Slavic example). On the other hand F_ST_ distance calculations are more suitable for populations living at distant, isolated geo-locations, where genetic divergence by sequence evolution is determining. Between fully isolated populations with no admixture, thus no shared mt Hgs, SHD values are always 1 regardless of their mutational divergence (sequence evolution), though their lineages can be distantly related. In this respect the two methods complement each other.

In sequence based genetic distance calculations each nucleotide counts equal, which may again lead to unrealistic inferences, as the fixation time between parent/leaf mt Hgs varies to a great extent between different mt Hgs. For example, there are 82 progeny lineages of major mt Hg H, all of which have just one mutation distance from their parent mt Hg, but their fixation time varies between 1735 and 12475 years [6]. Consequently considering each mutation distances equal is an approximation that is only statistically true leading to lower resolution and incoherencies especially at small genetic distances. In SHD calculation only the frequency of overlapping mt Hgs matters regardless of their age and other attributes. The only premise in SHD calculation is that shared mt Hgs have common origin, which link population histories in an extent proportional to the SHD value. Furthermore while mathematically F_ST_ is not a metric distance as it does not satisfy the triangle inequality, the SHD calculation complies with all metric axioms. This leads to consistent topology of hierarchic clustering based on the SHD matrix and also allows the global minimal distance optimization used in the MITOMIX calculations.

Considering the spatial distribution of populations and their multiple admixtures, the history and relations of most populations cannot be represented with a simplified bifurcating tree. Accordingly, while SHD calculation and hierarchical clustering can group similar populations to sub-branches, the resulting “phylogenetic” trees cannot visualize all the fine details of the underlying genetic connections between populations. This is especially true for migrating populations that share genetic elements with their ancestors, but subsequent admixtures may significantly alter their genetic pool. Identification of the admixing source from offspring generations – especially for diverse modern populations – is hindered by the numerous potential candidates carrying the same genetic components. Not surprisingly F_ST_ based mtDNA analysis indicated substantial genetic homogeneity of European populations [36], with only a few geographic or cultural isolates appearing to be genetic isolates as well [37]. On the other hand, studies of the Y chromosome [38, 39] and of autosomal diversity [40] revealed a general gradient of genetic similarity resulting from the complex admixture history of Eurasian populations [27, 41].

Our novel algorithm (MITOMIX) enables a hypothesis independent search for the best linear admixture combination (*ε*_1_, …,*ε_k_*) of *k* populations in order to minimize the SHD value from a test population. Since MITOMIX is based only on the composition of the mt Hgs regardless of their origin and age, it can be applied to any populations including neighboring and distant (migrating) ones. Concurring with the results of previous Y chromosome and genomic studies, MITOMIX analysis exposed a great extent of local and global admixtures. MITOMIX suggested the hypothetical composition(s) of the studied populations that provides insight into the subtle genetic connections, which alone cannot be studied by paired-distance measurements.

Fixation and spread of new Hg lineages takes considerable time and many generations. From the Hg frequency of available archaic and modern samples we demonstrated that in about ~8000 years the mtDNA phylogenetic tree has evolved just about 2 Hg sub-branches. Therefore at small to moderate time scales the mt tree evolution is insignificant, which is an ideal condition for applying the SHD calculation and MITOMIX methods. Besides, using an iterative method, we could successfully correct the mt Hg distribution shift of modern populations, resulting in a similar mt Hg distribution to that of their ancient ancestors (Fig. 2). This correction also considers sequence evolution and extends the applicability of the method to a larger time window. As we have shown, analysis of modern and archaic populations with corrected mt Hg data gives concordant results with previous studies testifying the applicability of the method. In case representative mitogenome data will be available from a series of historical time periods, MITOMIX can potentially identify all traces and the extent of admixture and uncover even as complex ethnogenesis as that of the European populations. Furthermore, in depth analysis of the age distribution of shared mt Hgs may provide a clue as to the periods of admixtures.

We also sequenced 272 modern Hungarian mtDNA genomes, significantly expanding the number of 17 publicly available Hungarian full-length mitogenomes. Applying low resolution genetic analyses Modern Hungarians were indicated to be genetically very similar to their European neighbors [42–44]. Our novel analysis not only confirmed those results but also revealed unpredicted finer details. Our SHD and MITOMIX analysis suggest that the majority of modern Hungarian maternal gene pool has ancient European origin, and their genetic history is surprisingly shared in a large extent with modern Danish, Belgian/Dutch, and Basque populations. A large part of this genetic similarity was definitely obtained in prehistoric Neolith-Bronze Age periods, but subsequent Iron-Age admixtures are also possible, as a significant fraction of the shared mt Hgs between modern Danes and Hungarians originated/expanded after the Bronze Age (Fig. 7). Genomic admixture data also supports this view, as a significant Northern European component was detected in modern Hungarian genomes [27], while the analysis of ancient European mitogenomes suggested that late Neolithic cultures had a key role in shaping modern Central European genetic diversity [45]. MITOMIX analysis indicated that a significant fraction of Hungarian mitogenomes are potentially originated from the Bell Beaker, Baltic Late Bronze Age and European Late Neolith/Bronze Age cultures, while a smaller portion came from admixture with neighboring Serbians, Croatians, Slovakians, Romanians, as well as other ancient populations (Northern Neolith Bronze Age, Hungarian Conquerors, Srubnaya cultures). Modern Hungarians identify themselves as having originated from the Hungarian Conquerors, whom are deemed to have brought Hungarians to the Carpathian Basin in the IXth century, leading to the establishment of the Kingdom of Hungary around 1,000. It is remarkable that only a minor genetic contribution (<3%) was detected from the analyzed Hungarian Conquerors, that is in line with other data indicating that Modern Hungarians contain only about 3-5% East Eurasian components [44, 46]. Our results suggest that either the available samples do not represent the entire Conqueror population or their number (and hence their genetic effect) was smaller than formerly estimated.

Recently, an increasing number of ancient mitogenome and genome sequences are being published as enrichment and NGS sequencing techniques are improving and becoming less expensive. However sequencing full genomes from degraded ancient samples is still challenging and costly, while isolating and sequencing mitogenomes is far easier. Given sufficient amount of mtHg data MITOMIX offers an affordable substitute to whole genome analysis. We also argue that while genomic admixture analysis results in much more information per sample, it involves an increased risk of sampling bias in population analysis because usually just a handful of samples are used to represent a population. It is a matter of course that our approach can be also used to analyze other non-recombinant hypervariable haploid loci such as chloroplast DNA and Y chromosome haplogroups.

The resolution of our and any other population genetic method largely depends on database quality and quantity. With increasing number of (sub)population-specific mitogenomes, the resolution of our SHD-based approaches will provide even more accurate results, which will help resolve some of the potentially incorrect relations. To facilitate this process, we provide our Eurasian mtDNA database in Supplementary Table 1, which surely necessitates further development. We encourage the mtDNA community to participate in the joint effort to create and publicly share a combined mitogenome database and also to undertake careful indication of the geographic and ethnic origin of all future sequences, independent of the original research objectives.

## Conclusions

SHD is a population genetic distance that complies with all metric axioms, which allows global minimal distance optimization used in MITOMIX. In case of closely related and/or admixing populations, SHD gives more realistic results and provides better resolution than F_ST_. MITOMIX can potentially identify all traces and the extent of admixture, offering an affordable substitute to whole genome admixture analysis.

Our results suggest that the majority of modern Hungarian maternal lineages have Late Neolith/Bronze Age European origins, and a smaller fraction originates from surrounding populations while only a minor genetic contribution (<3%) was identified from the IX^th^ Hungarian Conquerors. Our analysis shows that SHD and MITOMIX can augment previous methods by providing novel insights into past population processes.

## Methods

### mtDNA genotyping of modern Hungarian samples

We retrospectively analyzed 229 mitochondrial genomes (Illumina TruSight One) from our DNA bank and 43 previously published whole exome sequences [47]. The DNA samples were treated by the appropriate wet lab protocols recommended by the manufacturers. The trimmed, adapter removed paired-end reads (fastq) were mapped to the GRCh37 reference genome containing the rCRS (NC_012920) mtDNA reference sequence by the Burrows-Wheeler Aligner (version 0.7.9a-r786) [48] using the BWA MEM paired-end mapping algorithm. Duplicates were marked by the Picard tools (version 1.113) MarkDuplicates algorithm. The raw bam files were realigned and base quality recalibrated by Genom Analysis Tool Kit (GATK version: 3.3-0- g37228af) [49]. Individual sequences were compared to the rCRS (NC_012920) mtDNA reference sequence and classified into mitochondrial haplogroups by HaploFind [50].

### F_ST_ calculation

We applied the traditional sequence based method calculating pair-wise population differentiation values (Fst) with Arlequin 3.5.2.2 [51] from entire mtDNA genomes assuming a Tamura & Nei substitution model [52] with a gamma value of 0.325. Significant variations in Fst values were tested by 10,000 permutations between populations. As individual insertions and deletions make the alignment of multiple mtDNA genomes troublesome, only variable positions were aligned, and insertions and deletions were recoded to SNP-s as follows. Whole mtDNA genome fasta files were aligned to the NC_012920 human mtDNA reference sequence by an IUPAC code aware in-house aligner using the Needleman–Wunsch algorithm with weight parameters: match 6, IUPAC2match (R, Y, M, W, S, K) 3, IUPAC3match (B, D, H, V) 2, IUPAC4match (N) 1, mismatch −12, gap open −24, gap extend −6. Modern sequences with more than 500 missing or uncertain nucleotides (nt.) were excluded from further analysis. Then all nt. positions where any variation was detected were outputted to VCF files. Since Arlequin cannot manage VCF files SNPs, deletions and insertions were recoded by the following rules: nt-s with no variation at the given position were coded as the reference nt.; SNPs with variation were coded as the alternate allele; all insertions were coded as additional nt. letters, C for samples with reference sequence and T for samples containing the insertion; all deletions were also coded as additional nt. letters, T for samples with reference sequence and C for samples containing the deletion. Then Arlequin input files (arp) were generated from the recoded DNA sequences.

### Data visualization

The mt Hg age distribution of populations was visualized by violin plots using the ggplot2 (version 2.2.1) R (version 3.4.1) package [53].

The population distance matrixes (Slatkin’s linearized F_ST_ [30], and SHD) were clustered using hclust R (version 3.4.1) function with the ward.D2 linkage method [54, 55]. The hierarchic clusters were visualized by the ape R package (version 4.1) [56].

## Authors’ contributions

Z.M., T.K. and T.T. conceptualized SHD algorithm. M.M. provided mathematical proofs of SHD. Z.M. conceptualized MITOMIX algorithm. M.M., T.V. and Z.M. provided mathematical formalization of SHD, MITOMIX, and haplogroup frequency shift calculations. Z.M. implemented the SHD and MITOMIX software. P.B. and D.T. performed NGS wet-lab experiments. T.T., E.N. and Z.M. assembled the mitochondrial population database. Z.M. performed all formal analysis. T.T., T.K. and Z.M. evaluated experimental data. Z.M. and T.K. wrote the initial version of the manuscript while T.T. contributed immensely and I.R., I.N., Z.B., M.M., T.V., P.B. contributed to subsequent versions. All authors reviewed and approved the final version of the manuscript.

